# Simulating the evolutionary trajectories of metabolic pathways for insect symbionts in the *Sodalis* genus

**DOI:** 10.1101/819946

**Authors:** Rebecca J. Hall, Stephen Thorpe, Gavin H. Thomas, A. Jamie Wood

## Abstract

Insect-bacterial symbioses are ubiquitous, but there is still much to uncover about how these relationships establish, persist and evolve. The tsetse endosymbiont *Sodalis glossinidius* displays intriguing metabolic adaptations to its microenvironment, but the process by which this relationship evolved remains to be elucidated. The recent chance discovery of the free-living species of the *Sodalis* genus, *S. praecaptivus*, provides a serendipitous starting point from which to investigate the evolution of this symbiosis. Here, we present a flux balance model for *S. praecaptivus*. Metabolic modelling is used in combination with a multi-objective evolutionary algorithm to explore the trajectories that *S. glossinidius* may have undertaken after becoming internalised. The time-dependent loss of key genes is shown to influence the evolved populations, providing possible targets for future *in vitro* genetic manipulation. This method provides an unusually detailed perspective on possible evolutionary trajectories for *S. glossinidius* in this fundamental process of evolutionary and ecological change.

## 2 Introduction

Symbioses are both fundamental and ubiquitous in nature. Understanding their evolution poses an ongoing challenge, as well as an expanse of unresolved research questions. Bacterial symbionts of insects provide a range of benefits including stress tolerance (Dunbar et al., 2007; Wilcox et al., 2003), protection from predation (Nakabachi et al., 2013; Oliver et al., 2003; Wilcox et al., 2003) and the provision of metabolites (Aksoy, 1995; Hrusa et al., 2015; Manzano-Marín et al., 2015; Shigenobu et al., 2000; Snyder and Rio, 2015; Thomas et al., 2009). The latter forms arguably the strongest link within the symbioses. Host and symbiont frequently share metabolic substrates, as well as the products and components of individual biosynthetic pathways (McCutcheon et al., 2009a,b; McCutcheon and Moran, 2007; McCutcheon and von Dohlen, 2011; Thomas et al., 2009; Wilson et al., 2010). These relationships typically enable the host to survive on a nutritionally restricted diet, such as the blood meal of the tsetse (Michalkova et al., 2014; Rio et al., 2003; Snyder and Rio, 2015) or the plant sap that feeds the aphid (Akman Gündüz and Douglas, 2009; Baumann et al., 1995; Richards et al., 2010).

Deciphering the evolutionary pressures that affect the organisms within a symbiosis is an essential part of understanding the relationship. This includes establishing how the symbioses develop over time and the way in which the metabolism of the individuals is intertwined. It is, however, often hindered by biological difficulties. Symbiotic bacteria undergo genomic streamlining, may not be cultivatable *in vitro*, may no longer express stress response genes and might lack a sound outer membrane (Akman et al., 2002; Aksoy, 1995; Dale and Maudlin, 1999; Moran et al., 2008; Nakabachi et al., 2013; Pérez-Brocal et al., 2006; Wu et al., 2004). It is therefore impossible in many cases to test hypotheses about host-symbiont interactions in controlled experimental conditions. In these circumstances, computational techniques offer a viable, and currently the only, alternative to investigating metabolic potential and pseudogenisation in symbiotic bacteria.

Computational biology is now well established as a key tool of scientific discovery, now that vast amounts of data are generated quickly and cheaply from advancements in sequencing technology (Edwards et al., 2002; Kauffman et al., 2003). Genome scale metabolic modelling of microorganisms enables predictions to be made about metabolite preferences, transporter use and the functionality of biosynthetic pathways (Edwards et al., 2002; Lewis et al., 2012). Microbial metabolism can be simulated using flux balance analysis (FBA), a constraint-based quantitative approach that reconstructs a metabolic network from a genome annotation (Kauffman et al., 2003; Orth et al., 2010). FBA is a powerful tool when based on a well annotated genome and with the provision of *in vitro* experimental validation (Orth et al., 2010).

FBA is widely used for biotechnology applications, and this can be re-purposed to examine symbiosis. There are several published examples of using FBA to analyse the metabolism of symbiotic bacteria, including for *Buchnera aphidicola* (Macdonald et al., 2012, 2011; Thomas et al., 2009), *Sodalis glossinidius* (Belda et al., 2012; Hall et al., 2019), *Portiera aleyrodidarum* (Ankrah et al., 2017), *Hamiltonella defensa* (Ankrah et al., 2017) and strains of *Blattabacterium* (González-Domenech et al., 2012). There are also models published for the *Synechocystis* species used in the study of artificially induced symbiosis (Knoop et al., 2013, 2010; Nogales et al., 2012; Shastri and Morgan, 2005; Sørensen et al., 2016). FBA is useful in this instance, as experiments that would not be possible empirically, due to culturability issues, can be performed *in silico*. Furthermore, the genomes of symbiotic bacteria are often unusual, with large pathway deletions or widespread pseudogenisation (Dale and Maudlin, 1999; Goodhead et al., 2018; Shigenobu et al., 2000; Toh et al., 2006). Analysis of the resulting metabolic network via FBA can suggest whether these biosynthetic pathways are active, and predict which external metabolites might be required to support growth *in vitro*.

FBA has been applied to several microbiological problems. Boolean logical operators have been incorporated into *Escherichia coli* metabolic models to investigate the impact of gene regulation on a system (Covert and Palsson, 2002, 2003; Covert et al., 2001; Lee et al., 2006). Dynamic FBA, where a rate of change in flux constraints is included, has successfully modelled diauxic growth in *E. coli* (Mahadevan et al., 2002). FBA has been used to compare strains of *Blattabacterium* from separate cockroach lineages to assess their divergence (González-Domenech et al., 2012), and to predict the evolution of metabolism from *E. coli* experimental data sets (Harcombe et al., 2013). The evolution of metabolic networks in isolation has also been simulated with the aim of identifying key metabolites (Pfeiffer et al., 2005). FBA has not yet been harnessed to its full potential with regards to the investigation of symbiont evolution. This is perhaps surprising given that several models of *E. coli* metabolism are available as an evolutionary starting point (Edwards and Palsson, 2000; Feist et al., 2007; Orth et al., 2011; Orth and Palsson, 2012; Pál et al., 2006; Reed et al., 2003). The evolution of *B. aphidicola* and *Wigglesworthia glossinidia* from an *E. coli* ancestor has been simulated using FBA (Pál et al., 2006). This work, whilst elegant, has a key limitation. Reactions that are lost at the start have no chance of being reintroduced. This limits the evolutionary space that can be explored, as the loss of a key reaction at the start will fundamentally affect which reactions can be lost subsequently. A similar approach to that used by Pál et al. (2006) was used with dynamic FBA to study the evolution of cooperation and cross-feeding in *E. coli* (McNally and Borenstein, 2017). Using FBA in isolation to remove reactions successively may not therefore be the optimal way to simulate the evolution of symbiosis.

*In silico* evolution has been used increasingly in recent years to complement *in vivo* experimental evolution (Hindré et al., 2012). *In silico* evolution benefits from being able to test widely different ecological conditions whilst controlling key variables (Batut et al., 2013). For example, it allows the investigation of groups of mutations that lead to a specific phenotype, or mutations that are difficult to induce *in vitro* (François and Hakim, 2004). This has enabled the study of many aspects of evolution, including simulating the reduction of genome size in an individual (Batut et al., 2013). Multi-objective evolutionary algorithms (MOEA) have been used in many disciplines for solving problems that have two or more conflicting objectives (Budinich et al., 2017). The use of MOEA in combination with metabolic models has been implemented for the design of minimal genomes (Wang and Maranas, 2018) and for the production of industrially relevant molecules (Fong et al., 2005; Garcia and Trinh, 2019). It has however seen only limited use for *in silico* evolution (Machado and Herrgård, 2015). When viewed computationally, the evolution of symbiosis can be considered as a multi-objective optimisation; symbiotic bacteria undergo genome reduction whilst trying to maximise their individual growth.

A free-living organism within the *Sodalis* genus has been characterised and sequenced recently (Chari et al., 2015; Clayton et al., 2012). *Sodalis praecaptivus* was isolated from a human wound, caused by an impalement on a crab apple tree branch. It is assumed that the tree was the likely source of the *S. praecaptivus* infection. *S. praecaptivus* is a prototroph, capable of growth in minimal media and at 37°C (Chari et al., 2015). The annotated genome sequence for *S. praecaptivus* is also available (Clayton et al., 2012). It is of particular interest given its close relation *Sodalis glossinidius*, secondary symbiont of the tsetse (Dale and Maudlin, 1999). The tsetse, genus *Glossina* is medically important as the vector for *Trypanosoma brucei*, causative agent of human African trypanosomiasis (WHO, 1998). *S. praecaptivus* therefore provides a rich set of data from which to begin investigations into the origin of, and adaptations within, the tsetse-*S. glossinidius* symbiosis.

Here, we present a flux balance model for *S. praecaptivus, i* RH830. This model, and a model of *S. glossinidius* metabolism, *i*LF517 (Hall et al., 2019), both represent adaptations of the organisms to their contrasting environments. The *Sodalis* system is therefore an excellent candidate for assessing the ability of FBA to describe the evolution of symbioses. A MOEA has been used to evolve *i*RH830 under various biological conditions. The aim was to investigate computationally the route that *S. glossinidius* may have taken in its transition to symbiosis. It is not known whether the solutions found by *S. glossinidius*, described in *i*LF517, are the only possible outcomes given the metabolic constraints of the microenvironment, or whether the symbiont’s unusual metabolic network evolved by chance. The application of the MOEA to *i*RH830 enabled the observation that certain key pseudogenisations may have occurred earlier in the symbiosis than previously thought. The effect of exposing the ancestral *Sodalis* to contrasting diets was also modelled, mirroring the different trajectories that this genus has taken within blood- and sap-feeding insects. It is hoped that the techniques used here can be applied to other symbiotic systems to drive forward the discovery of novel relationship criteria.

## 3 Results

### 3.1 A model of *S. praecaptivus* metabolism, *i*RH830

In order to investigate computationally the path that *S. glossinidius* has taken to symbiosis, a metabolic model describing its close, free-living relative *S. praecaptivus* was constructed (Figure 1). Full details are given in Supplementary File 1. *i*RH830 is contains 830 genes, 891 metabolites and 1246 reactions (excluding pseudoreactions), and is a prototroph for all essential amino acids. An iterative process of gap filling was undertaken by comparing the draft *S. praecaptivus* model to *i*LF517 (*S. glossinidius*) and *i*JO1366, a model of *E. coli* metabolism (Orth et al., 2011). *i*RH830 is supplied with an oxygen uptake value of 20 mmol gr DW^-1^ hr^-1^, reflecting the highly aerated conditions the organism is grown in and to retain consistency with models of *E. coli* metabolism (Orth et al., 2011; Reed et al., 2003).

**Figure 1:**
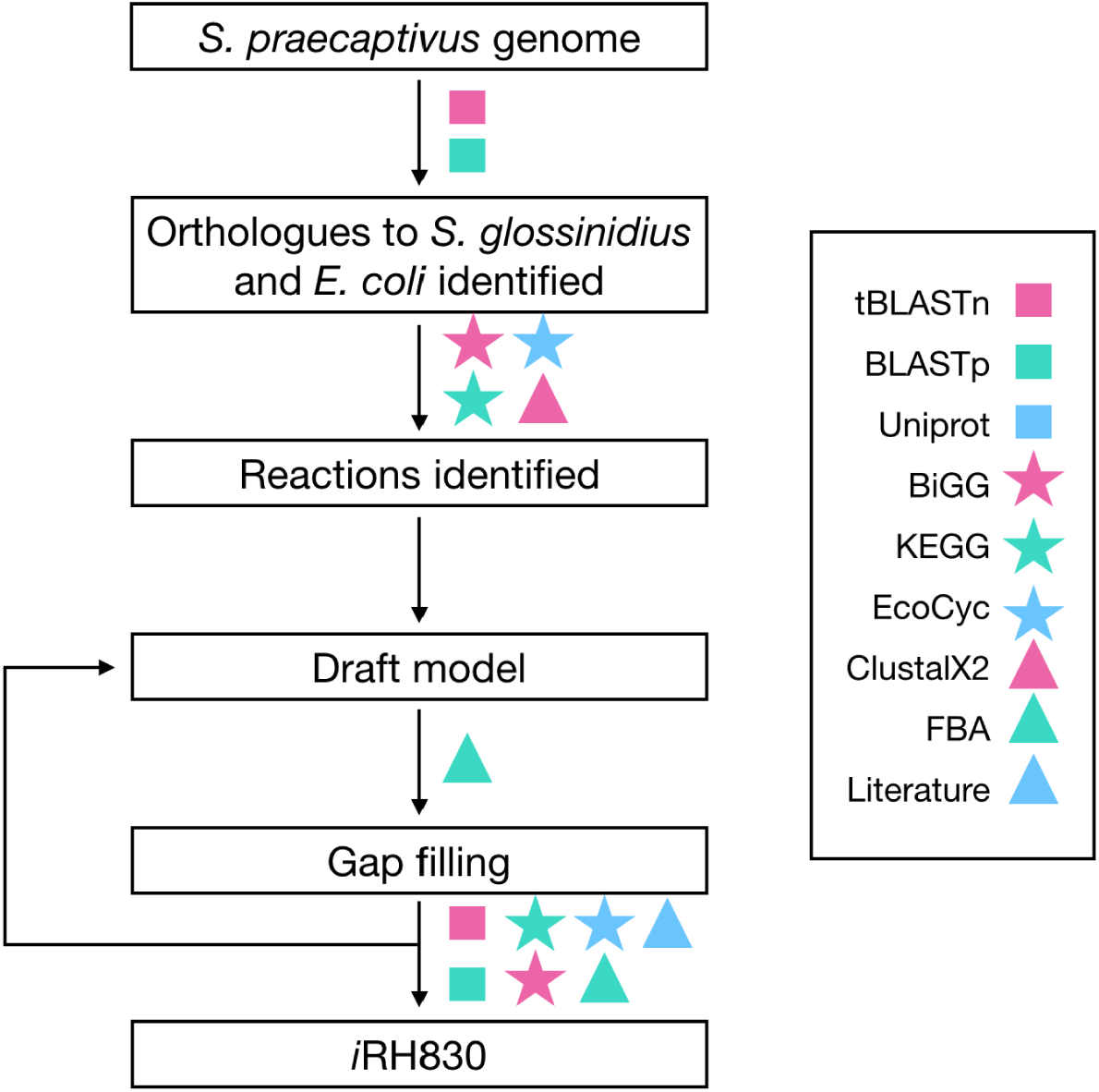
The construction process for *i*RH830. The *S. praecaptivus* genome was mined for orthologues to metabolic genes in *E. coli* and *S. glossinidius*, before compiling into a draft model. An iterative process of testing and gap filling was then performed, using information provided in various databases.

### 3.2 *S. praecaptivus* can grow on the unusual sugar alcohol xylitol

Computational models are strengthened when accompanied by experimental verification of their *in silico* predictions. A series of biochemical screens was therefore conducted using Biolog phenotypic microplates to investigate carbon utilization by *S. praecaptivus*. In total, 190 metabolites were tested for their ability to act as the sole carbon source for *S. praecaptivus*. Experiments were conducted in triplicate with full results detailed in Supplementary File 2. Through this phenotypic screen, it was found that *S. praecaptivus* was able to use 19 of the metabolites tested as a sole source of carbon (Table 1). When these metabolites were tested *in silico* by the exogenous addition to *i*RH830, it was found that all but two mirrored the *in vitro* data qualitatively; *N*-acetyl D-galactosamine (GalNAc) and xylitol. GalNAc did not produce a viable biomass output, and xylitol was not included as a metabolite in *i*RH830 initially. Xylitol is not found in any prokaryotic model in the BiGG Models database, hence it had not been considered for inclusion in the construction of *i*RH830.

**Table 1:**
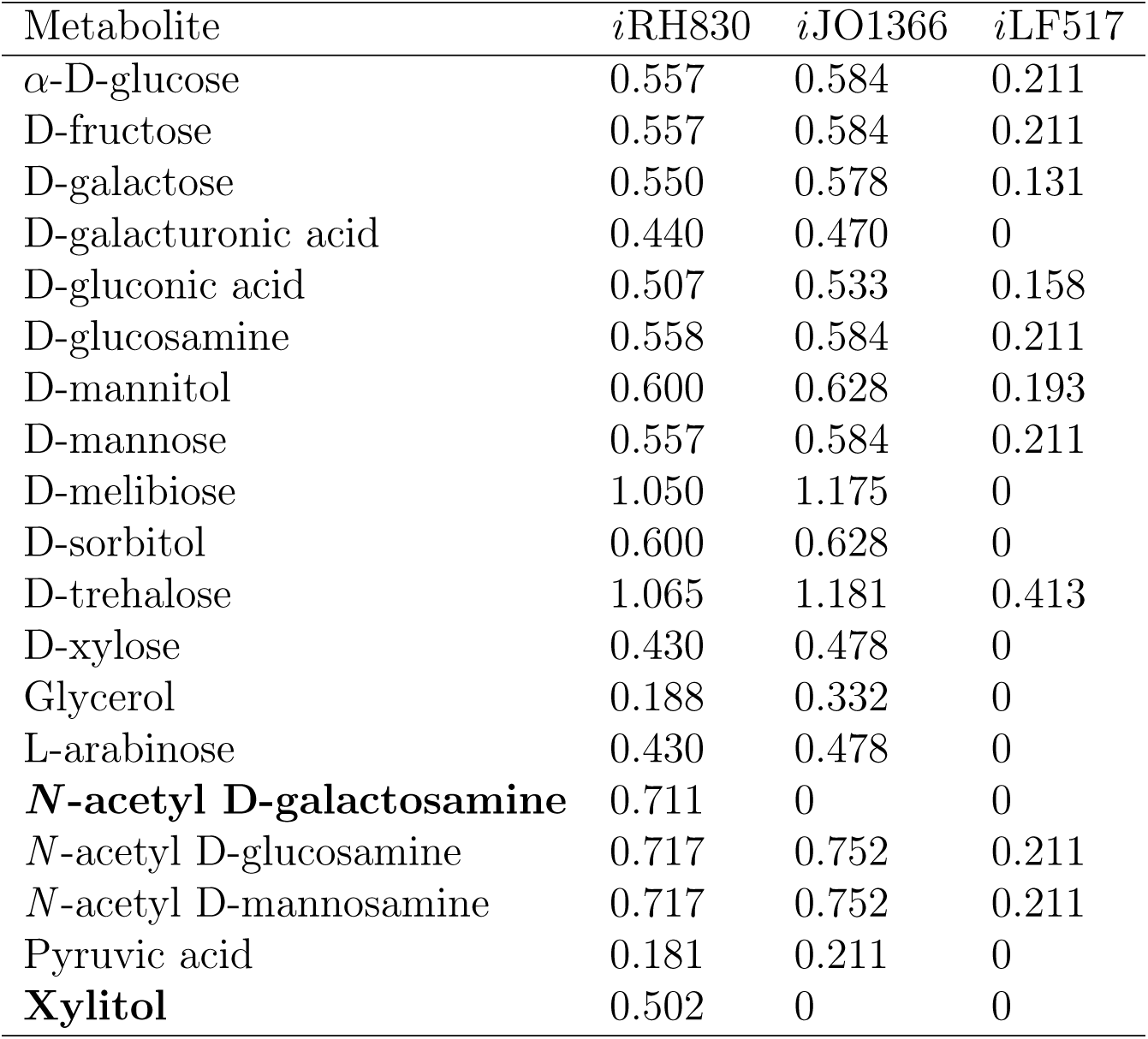
Positive results from one representative *S. praecaptivus* phenotypic screen. *i*RH830, *i*JO1366, and *i*LF517 ?biomass outputs (gr DW (mmol glucose)^-1^ hr^-1^) following the addition of these metabolites to models currently not supplied with a sugar. Metabolites in **bold** initially did not produce a positive biomass output in early iterations of the *S. praecaptivus* model.

| Metabolite | *i*RH830 | *i*JO1366 | *i*LF517 |
| --- | --- | --- | --- |
| α-D-glucose | 0.557 | 0.584 | 0.211 |
| D-fructose | 0.557 | 0.584 | 0.211 |
| D-galactose | 0.550 | 0.578 | 0.131 |
| D-galacturonic acid | 0.440 | 0.470 | 0 |
| D-gluconic acid | 0.507 | 0.533 | 0.158 |
| D-glucosamine | 0.558 | 0.584 | 0.211 |
| D-mannitol | 0.600 | 0.628 | 0.193 |
| D-mannose | 0.557 | 0.584 | 0.211 |
| D-melibiose | 1.050 | 1.175 | 0 |
| D-sorbitol | 0.600 | 0.628 | 0 |
| D-trehalose | 1.065 | 1.181 | 0.413 |
| D-xylose | 0.430 | 0.478 | 0 |
| Glycerol | 0.188 | 0.332 | 0 |
| L-arabinose | 0.430 | 0.478 | 0 |
| **N-acetyl D-galactosamine** | 0.711 | 0 | 0 |
| N-acetyl D-glucosamine | 0.717 | 0.752 | 0.211 |
| N-acetyl D-mannosamine | 0.717 | 0.752 | 0.211 |
| Pyruvic acid | 0.181 | 0.211 | 0 |
| **Xylitol** | 0.502 | 0 | 0 |

To investigate the use of xylitol as a sole carbon source by *S. praecaptivus*, cultures were established in a 96-well microplate with xylitol supplemented into M9 minimal medium at concentrations of 12.5 mM, 5 mM, and 500 *µ*M. *S. praecaptivus* was able to use xylitol equally well at 12.5 mM and 5 mM concentration (an optical density of approximately 0.27 at 650 nm (OD_650_), with a small amount of growth observed at 500 *µ*M xylitol (a final OD_650_ of approximately 0.14) (Figure 2A). *S. praecaptivus* was also shown to reach an OD_650_ of approximately 0.39 after 36 hours when 25 mM GalNAc was added as the sole carbon source (Supplementary Figure S1). This confirms the qualitative observations of the large-scale screen.

**Figure 2:**
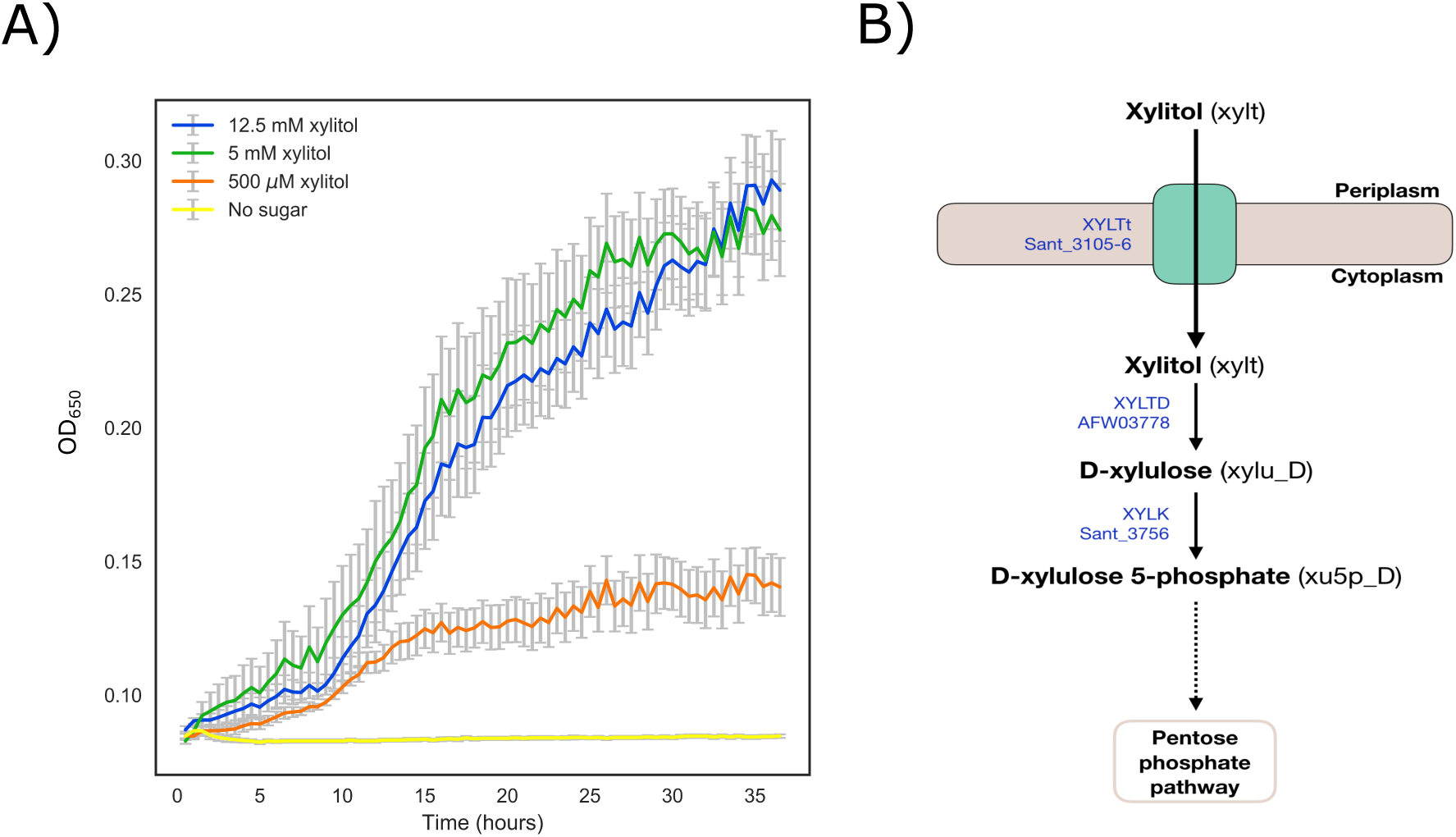
Xylitol use by *S. praecaptivus*. (**A**) Growth of *S. praecaptivus* on 12.5 mM (blue), 5 mM (green) and 500 *µ*M (orange) xylitol, and in M9 salts with no added sugar (yellow) over 36 hours. Measurements in sextuplicate, error bars SEM. (**B**) The putative xylitol degradation pathway in *S. praecaptivus*. Reaction names and *S. praecaptivus* gene assignments are given in blue.

KEGG was then used to identify organisms with xylitol degradation pathways. The sequence for a xylitol dehydrogenase from *Morganella morganii* (MU9_3130) was extracted from UniProt and used in a protein BLAST search against *S. praecaptivus*. From this, a hypothetical xylitol dehydrogenase was identified (NCBI reference AFW03778) with 79.7% sequence identify to MU9_3130 (Figure 2B). This reaction converts xylitol into D-xylulose. The reaction to convert D-xylulose into D-xylulose 5-phosphate, leading to the pentose phosphate pathway, was already included in the draft model (Sant_3756). The periplasmic binding protein L580_2330 from *Serratia fonticola* was used to aid the discovery of components of the xylitol transporter XltABC in *S. praecaptivus* (Sant_3104-6). These discoveries enabled the inclusion of two new reactions into the draft model, XYLTD (xylitol dehydrogenase) and XYLTt (putative xylitol transporter) (Figure 2B).

Neither models of *S. glossinidius* (*i*LF517) or *E. coli* (*i*JO1366) were able to produce a positive biomass output with xylitol as a sole carbon source (Table 1). To investigate whether these organisms, like *S. praecaptivus*, have the genomic capacity to metabolise xylitol, the *S. glossinidius* str. *morsitans* and *E. coli* str. K-12 substr. MG1655 genomes were mined using translated nucleotide BLAST searches with the *S. praecaptivus* proteins as queries. *S. glossinidius* appears to have retained all components of the xylitol transporter, xylitol dehydrogenase and xylulose kinase found in *S. praecaptivus* (Table 2). The *E. coli* genome, in contrast, does not contain xylitol degradation genes orthologous to *S. praecaptivus* (Table 2). There are candidate genes for all except Sant_3725, but most of these are of low similarity. Two show a higher percentage identity, a deaminase (82.61%) and a kinase (86.22%), possibly involved in the transport and metabolism of other, similar carbon sources. This indicates that species of the *Sodalis* genus, unlike *E. coli*, can import and metabolise xylitol.

**Table 2:** Candidate components of the xylitol metabolic pathway in *E. coli* and *S. glossinidius*. Candidate proteins based on tBLASTn searches using the *S. praecaptivus* proteins as search queries. Percentage identity to *S. praecaptivus* sequences, predicted number of amino acids (aa), locus tags (SGGMMB4…) and NCBI descriptions (*italics*) are presented. Top tBLASTn results are given.

| *S. praecaptivus* | *E. coli* K-12 substr. MG1655 | *S. glossinidius* |
| --- | --- | --- |
| **Sant_3104** | *RbsB* | SGGMMB4_01390 |
| 333 aa, SBP | 30.69%, 281 aa | 97.90%, 333 aa |
| **Sant_3105** | *D-ribose transporter* | SGGMMB4_01389 |
| 345 aa, permease | 41.64%, 308 aa | 92.17%, 345 aa |
| **Sant_3106** | *ATP binding protein* | SGGMMB4_01388 |
| 501 aa, ATP-binding | 44.02%, 491 aa | 96.81%, 501 aa |
| **Sant_3107** | *Xylulose kinase* | SGGMMB4_01387 |
| 504 aa, xylulose kinase | 29.70%, 483 aa | 89.29%, 504 aa |
| **AFW03778** | *Alcohol dehydrogenase* | SGGMMB4_01386 |
| 344 aa, putative dehydrogenase | 38.01%, 317 aa | 92.73%, 344 aa |
| **Sant_3756** | *Xylulokinase* | *Xylulose kinase* |
| 484 aa | 62.86%, 482 aa | 27.27%, 417 aa |

### 3.3 Robustness analysis of the *S. praecaptivus* metabolic network

Robustness analysis was used to examine reaction essentiality and therefore redundancy in the *i*RH830 network. *i*RH830 was run on a tsetse-specific nutrient limited medium (“famine”) and a blood medium simulating the internal tsetse environment and informed by *S. glossinidius* requirements (Hall et al., 2019) (“blood”, Table S1). All media is detailed in Supplementary File 1. Reactions were removed individually and the resulting effect on biomass output noted. The same analysis was also run on *i*LF517 in blood as a comparison.

There are 282 essential reactions in *i*RH830 when the medium (famine) is nutritionally limited, and 228 in the tsetse-specific blood medium (Figure 3A). The overall pattern for the two conditions is very similar. The subsystem most represented in either condition is for cofactor and prosthetic group biosynthesis, with 88 and 87 essential reactions for the famine and blood media, respectively. The main difference between blood and famine at the subsystem level can be attributed to amino acid metabolism. Of the 228 reactions essential in blood, 15.8% are involved in amino acid metabolism. In famine, which is not supplied with amino acids, 48 more essential reactions and a greater proportion of the total (29.8%) are used for amino acid metabolism. There are also three more essential transport reactions when the medium is limited.

**Figure 3:**
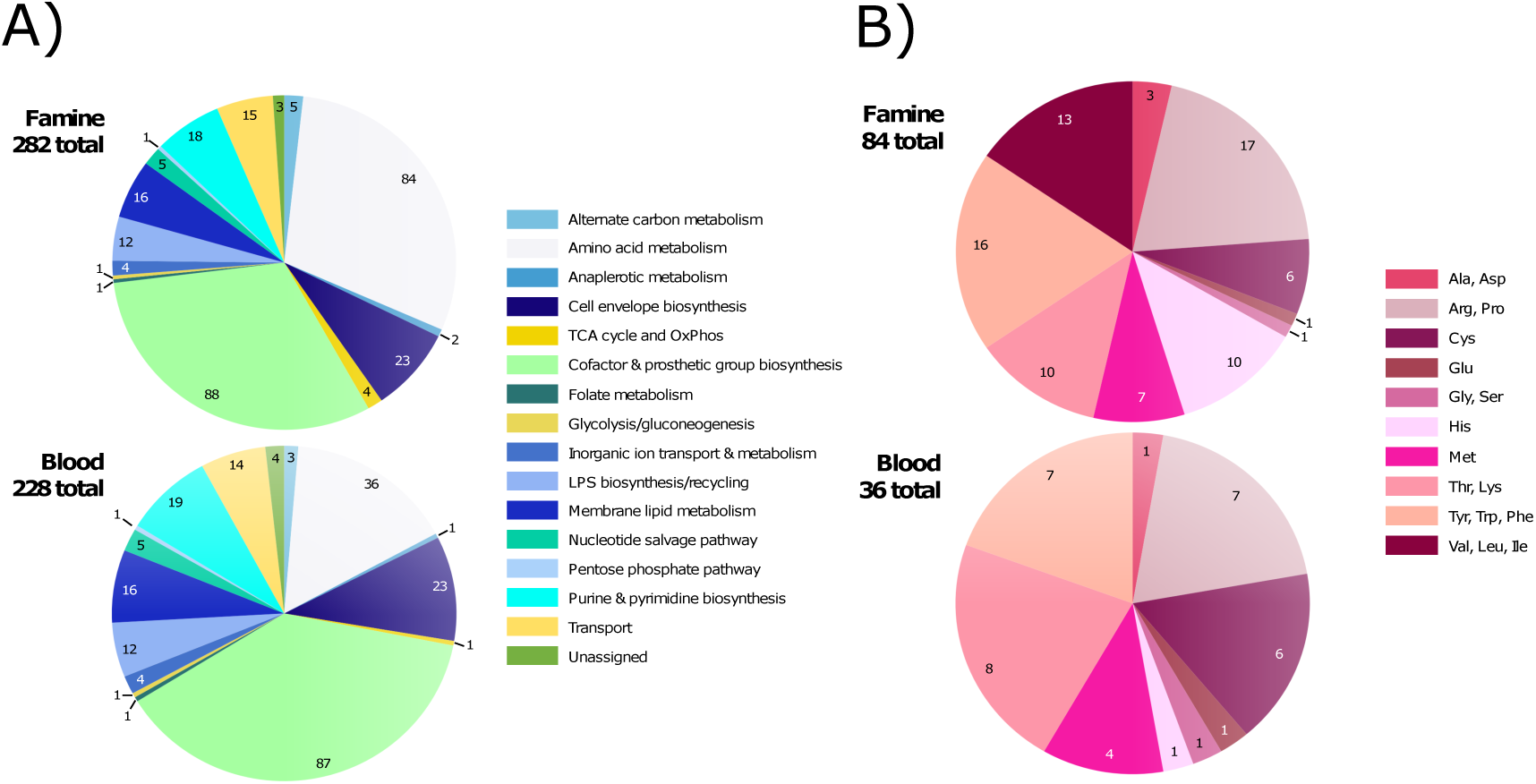
Robustness analysis of *i*RH830. (**A)** Essential reactions in famine (top) and blood (bottom) media. Essential reactions are categorised by subsystem. (**B**) Essential reactions involved in amino acid metabolism in *i*RH830 in famine (top) and blood (bottom) media.

The reactions involved in amino acid metabolism were then analysed further to ascertain whether the two conditions differ in composition as well as size. Of the 84 essential reactions in the famine medium, the highest number (17) are involved in the metabolism of L-arginine and L-proline, followed by L-threonine and L-lysine (16) (Figure 3B). There are 13 essential reaction for the metabolism of L-valine, L-leucine and L-isoleucine. There are two subsystems with the fewest essential reactions; L-glutamate (one), and glycine and L-serine metabolism (one). A single reaction in *i*RH830 (GHMT2r, glycine hydroxymethyltransferase) converts glycine to L-serine via a reversible reaction.

There are fewer essential reactions for amino acid metabolism when *i*RH830 is supplied with the blood medium, and the pattern of essentiality differs between the two media (Figure 3B). In blood, the two subsystems with the highest proportions mirror that seen in famine; L-threonine and L-lysine (eight), L-arginine and L-proline (seven). The pathways for L-tyrosine, L-tryptophan and L-phenylalanine also have seven essential reactions. The main difference is that there are no essential reactions involved in the metabolism of L-valine, L-leucine and L-isoleucine when *i*RH830 is supplied with a blood medium, compared to the 13 counted in famine. There is also only one reaction essential for L-histidine metabolism in blood, whereas there are 10 under the famine conditions. Overall, therefore, the main difference observed when *i*RH830 is supplied with contrasting metabolite availability lies in amino acid metabolism.

### 3.4 Media provisioning affects evolutionary trajectories

NSGA-II is a heuristic multi-objective optimisation algorithm used to evaluate multi-objective problems without giving weight to any specific outcome. Evolution within a constrained environment, such as the tsetse microenvironment, can be considered a multi-objective optimisation problem of trying to reduce the genome size to increase replication speed, while still retaining sufficient capacity to grow (Moran, 1996). The MOEA was used to explore the potential evolutionary trajectories of *S. praecaptivus* when exposed to similar environmental conditions to *S. glossinidius*. A graphical description of the MOEA is provided in Section 5.5. The conditions under which *i*RH830 and *i*LF517 were evolved are detailed in Table 3. In Scenario i, *i*RH830 was evolved in blood and famine growth media, as well as a medium that mimics plant sap (Supplementary File 1), to examine the effect of metabolite availability. In Scenario ii, three key reactions were removed from *i*RH830 prior to evolution to compare the trajectories that arise as a result of pseudogenisations. In Scenario iii, the MOEA is applied to *i*LF517 to investigate the possible future of *S. glossinidius* as a symbiont.

**Table 3:** *In silico* evolution conditions. Conditions under which *i*RH830 and *i*LF517 were evolved, including wild-type (WT) or reaction knockouts and media type.

| Scenario | Test | Model | Media | Section |
| --- | --- | --- | --- | --- |
| i | Effect of growth media | *i*RH830 (WT) | Blood, sap, famine | 3.4 |
| ii | Effect of gene loss | *i*RH830 (ΔASPTA, ΔPDH, ΔPPC) | Blood | 3.5 |
| iii | Future of *S. glossinidius* | *i*RH830 (WT), *i*LF517 (WT) | Blood | 3.6 |

Species of the *Sodalis* genus have been found in insects that feed on a variety of contrasting diets, including blood (e.g. tsetse (Dale and Maudlin, 1999) and ticks (Boyd et al., 2016; Chrudimský et al., 2012; Novakova and Hypsa, 2007)) and plant tissue (e.g. weevils (Oakeson et al., 2014)). To replicate *Sodalis* evolution in different environments, the MOEA was applied to *i*RH830 that was supplied with a tsetse-specific blood medium, a medium that mirrors plant sap, and a nutritionally limited “famine” medium (Table 3, Scenario i). The algorithm underwent ten runs of 3000 generations and the resulting solutions were collated.

In all conditions, the models evolved to completion, demonstrated by the convergence of solutions to the left of the plots (Figure 4A). The number of reactions decreases over evolutionary time, with the majority of solutions clustering at the maximum biomass output. This is an indication that sub-optimal solutions are being removed successfully. After 3000 generations, there are a range of solution sizes at the maximum biomass output found in sap, whereas in blood and famine all solutions at this time point cluster at the minimum number of reactions. The two rich media, blood and sap (Figure 4A), produce a lot of metabolic flexibility, with a complete range of possible biomass outputs produced by the smallest models. When grown in the nutritionally limited famine medium, there is significantly less flexibility in terms of possible solutions found (Figure 4A). Here, the majority of the solutions cluster at the minimum reactions/maximum biomass output. This is as expected, given the fitness function of the MOEA. In blood and sap, the biomass outputs reach near zero, made possible by the variety of available substrates. In famine, the options for streamlining are limited, resulting in few solutions that are able to deviate away from what is selected by the fitness function.

**Figure 4:**
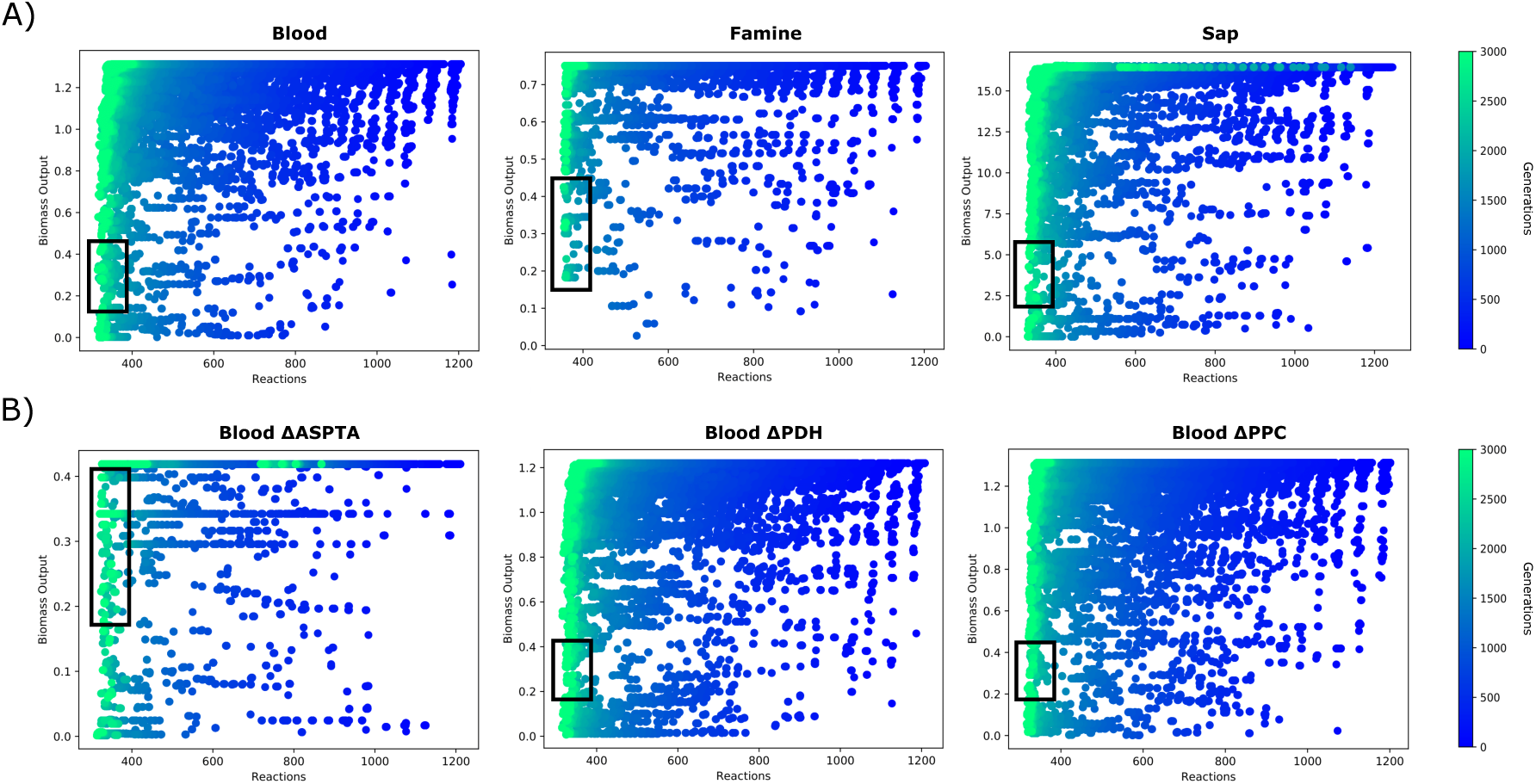
*i*RH830 evolved under different starting conditions. (**A)** A tsetse-species blood medium (left), a medium mimicking plant sap (centre) and a nutritionally limited famine medium (right). (**B**) *i*RH830 evolution in a blood medium with the reactions ASPTA (left), PDH (centre) and PPC (right) removed at the start. The MOEA was run for 3000 generations, with the plot depicting new populations every 50 generations (blue to green). Black boxes indicate individual solutions selected for further analysis.

A number of individual solutions that were representative of the biomass output of *i*LF517 (Hall et al., 2019) were then selected from each of these simulations (Figure 4, black boxes). The raw, binary data were translated back into reaction names and this was subsequently processed to produce a list of “core non-essential reactions”. These reactions are found in all individuals selected, and do not produce a lethal phenotype when removed. A full list of all core non-essential reactions described here can be found in Supplementary Table S2. There are 14 core non-essentials reactions found in all 1194 of the individuals examined when *i*RH830 was supplied with blood; AGDC, ARGabc, ASNt2r, G6PDA, H2Ot, HISt2r, ILEt2r, NH4t, RPE, TKT1, TKT2, TMK, TRPt2r and TYRt2r. In sap, only one non-essential reaction is found in all 1888 individuals; the L-arginine ABC transporter reaction ARGabc. As anticipated, when grown in the limited famine medium there are a higher number of core non-essential reactions (22 found in each of the 2989 individuals tested); ATPS4r, CO2t, ENO, FORt, GAPD, GHMT2r, GLUDy, ORNDC, PAPSR, PGCD, PGK, PGM, PPPGO3, PSERT, PSP_L, RPE, TALA, THRAi, TKT1, TPI, TRDR and TRPS1. The rare core non-essential reactions were then calculated. In famine, there are 73 unique reactions that occur in less than 0.1% of the 2989 evolved models. This is significantly more than for sap (13 in less than 0.1% of 1888 models) and blood (five in less than 0.1% of 1194 models).

These core non-essential reactions were then analysed by subsystem to assess themes across the different conditions. In blood, over half (eight of 14) of these are secondary transporter reactions (Figure 5A). This reflects what is observed in *S. glossinidius*, which has retained, for example, secondary amino acid transporters (Hall et al., 2019). The loss of metabolic pathways and the maintenance of functional transporters is characteristic of symbiotic bacteria that are able to scavenge metabolites from their microenvironment. As mentioned previously, the only core non-essential reaction in sap is a transport reaction. In contrast, the set of core non-essentials are more varied when metabolites are limited (famine), with a particular emphasis towards central metabolism and amino acid metabolism.

**Figure 5:**
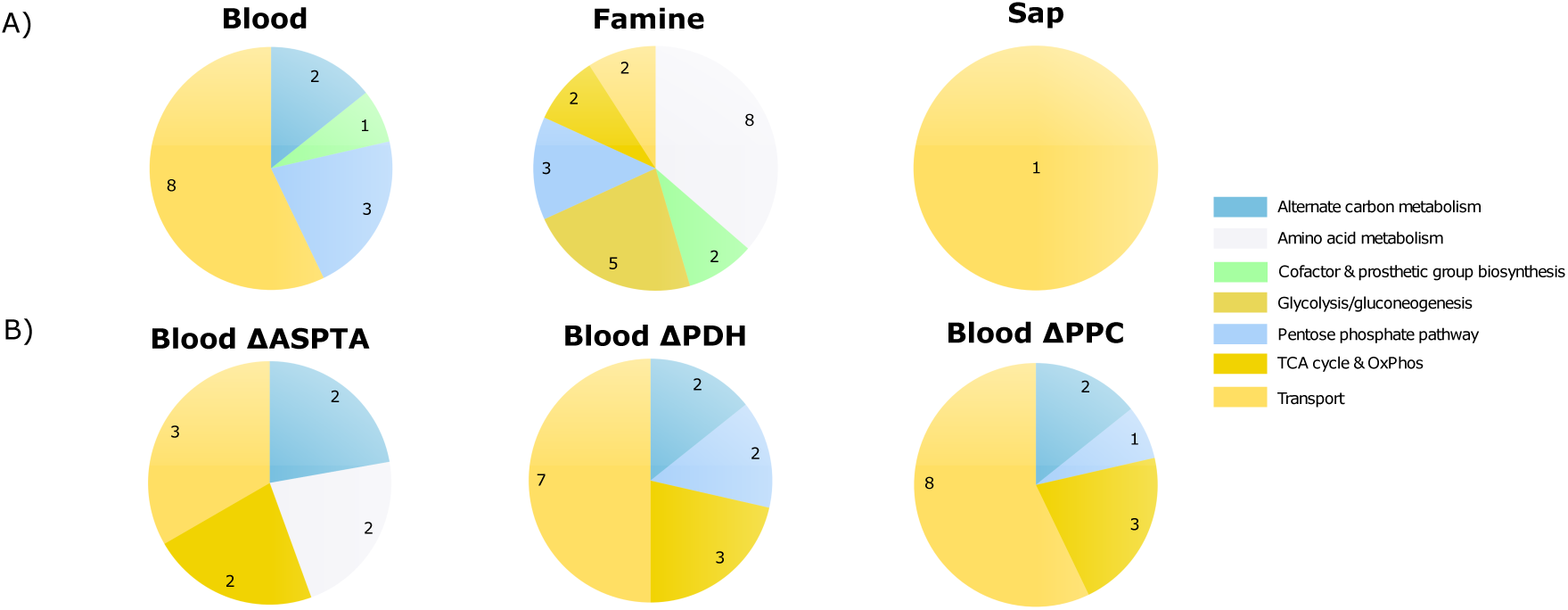
Core non-essential reactions in evolved *i*RH830 populations. (**A)** The proportion of core non-essential reactions per conditions by subsystem when the ancestral *i*RH830 is exposed to blood (left), famine (centre), or sap (right) media. (**B**) Core non-essential reactions in ΔASPTA (left), ΔPDH (centre), and ΔPPC (right) *i*RH830 models in a blood medium, grouped by subsystem.

**Figure 6:**
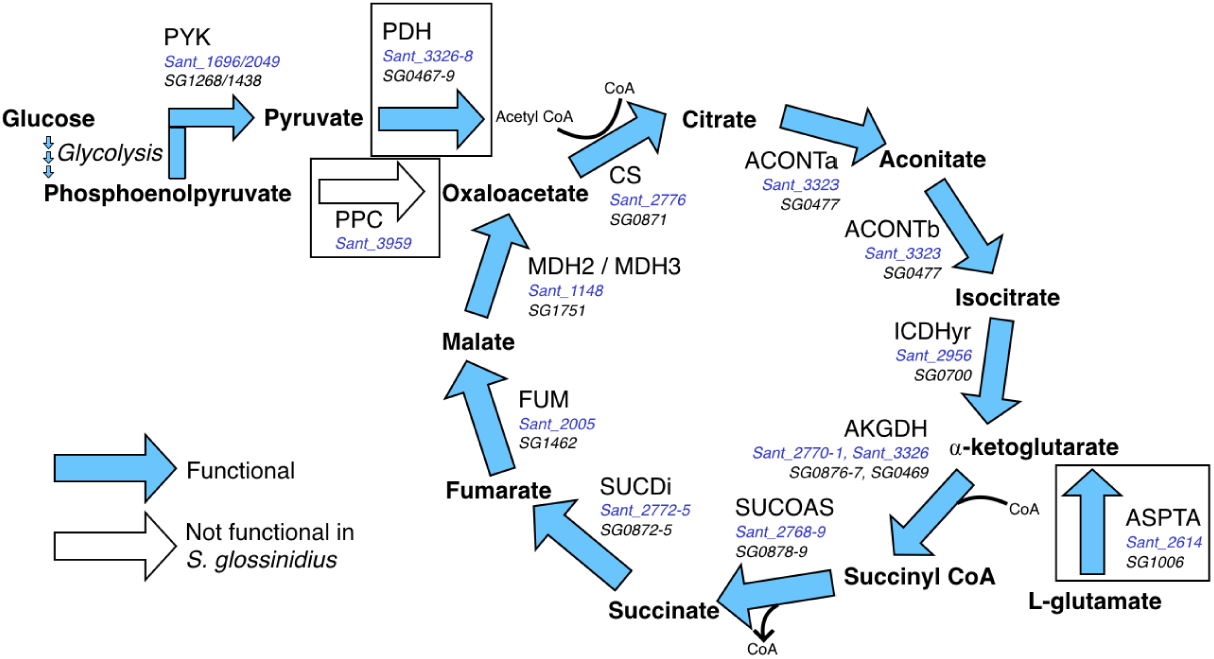
TCA cycle reactions examined by the MOEA. Three reactions were removed from *i*RH830 to investigate the resulting trajectories following application of the MOEA; ASPTA, PDH and PPC. Blue arrows show reactions functional in *S. praecaptivus* and *S. glossinidius*, white arrows show reaction not functional in *S. glossinidius*. Gene associations in *S. praecaptivus* and *S. glossinidius* are given in blue and black text, respectively. Adapted from Hall et al. (2019).

### 3.5 Time-dependent gene loss can be predicted

A characteristic of *S. glossinidius* and other symbiotic bacteria is their propensity to accumulate pseudogenes. It is not known whether key genes are lost early in the tsetse-*Sodalis* symbiosis to enable the initiation, or whether their loss is an inevitable consequence of genomic streamlining. To investigate the effect that pseudogenising key genes early in evolutionary time has on the trajectory of a symbiont, the MOEA was run on *i*RH830 with one of three reactions removed at the start, with the resulting solutions compared to wild-type (WT) (Table 3, Scenario ii). The reactions selected were PPC (phosphoenolpyruvate carboxylase), as a key pseudogenisation in *S. glossinidius* central metabolism (Hall et al., 2019), and PDH (pyruvate dehydrogenase) and ASPTA (aspartate transaminase) as two other reactions that feed the TCA cycle.

When considering the population plots, there is minimal qualitative difference between ΔPDH and ΔPPC (Figure 4B). ΔASPTA, in contrast, produces solutions with a much lower biomass output and with fewer individuals that deviate away from the optimum as defined by the fitness function. A selection of individuals were then selected and the number of core non-essential reactions in the evolved models were then analysed as described previously (Figure 4B, black boxes). There are one, 11 and nine core non-essential reactions in the WT, ΔPDH and ΔPPC solutions, respectively, whereas there are 61 in ΔASPTA. These 61 reactions function in a variety of subsystems, particularly transport, central metabolism, amino acid metabolism and nucleotide salvage pathways. There are minimal differences between the core non-essential reactions at the subsystem level between ΔPDH and ΔPPC (Figure 5B). The main difference of note is the presence of reactions involved in amino acid metabolism in the ΔASPTA, but not the ΔPDH or ΔPPC solutions. The time-dependent nature of pseudogenisations can therefore be estimated, using the resulting evolutionary trajectories as a guide.

### 3.6 A prediction of the evolutionary future of *S. glossinidius* as a symbiont

*S. glossinidius* is a secondary symbiont. Both bacterium and insect can survive independently of one another, and the former is likely a more recent acquisition. It is however unclear how recently *S. glossinidius* was captured, or, given the pseudogenisations already present, how much more streamlined its genome can become. The algorithm was therefore applied to *i*LF517 in a blood medium with the aim of evaluating potential future evolutionary trajectories within the bounds of its relationship with host and primary symbiont (Table 3, Scenario iii).

There are a spread of biomass outputs found at the end of the evolution (Figure 7A), as observed when *i*RH830 was evolved in blood. The smallest solutions contain approximately 300 reactions. The evolved solutions were then compared to the evolved *i*RH830 models to assess their similarity. Ten evolved models for both *i*RH830 and *i*LF517 were analysed. All exchange reactions and those that carried zero flux were removed from further analysis.

**Figure 7:**
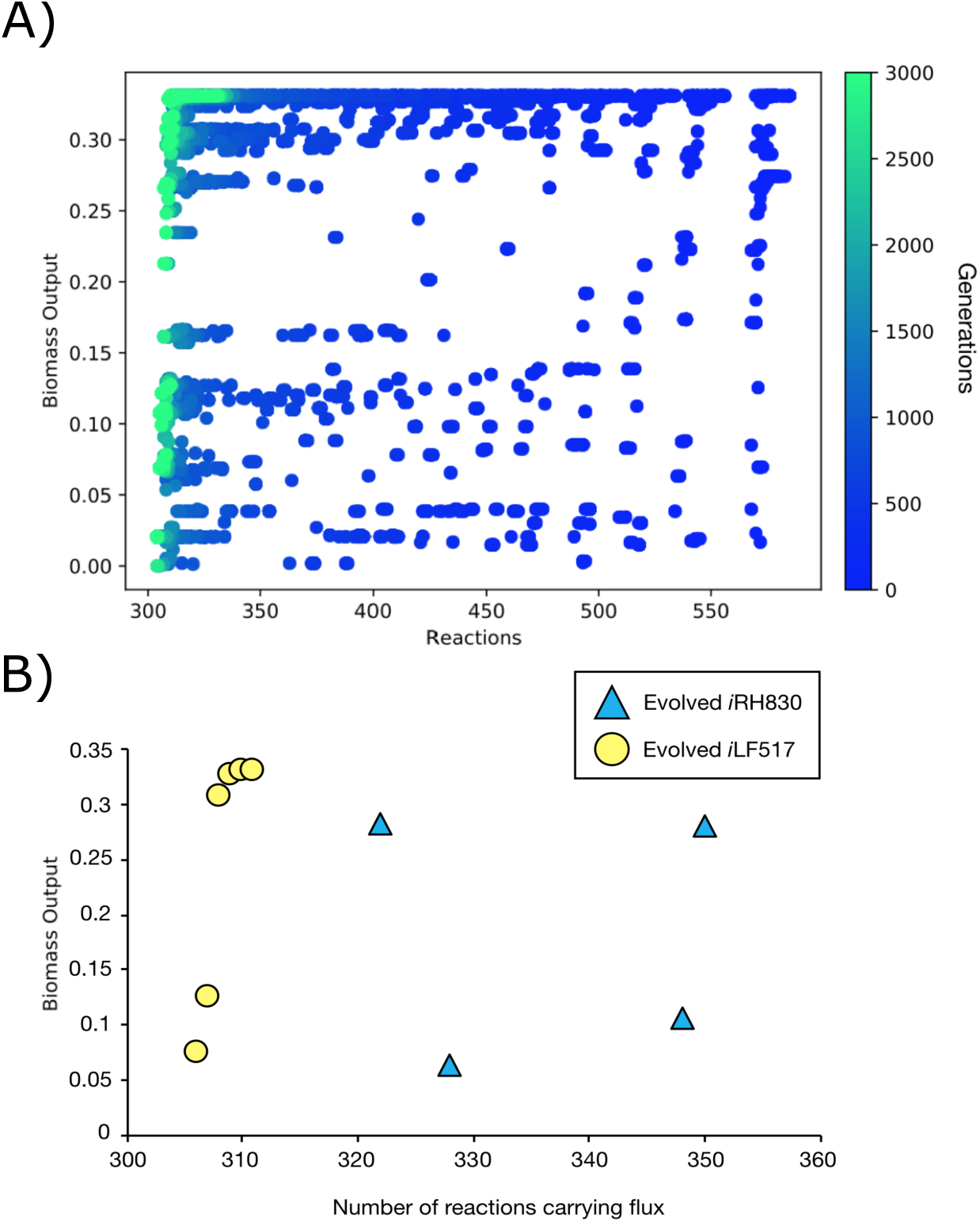
Evolution of *i*LF517. (**A)** Evolution of *i*LF517 in a blood medium. MOEA was run for 3000 generations (blue to green). (**B**) Biomass output (gr DW (mmol glucose)^-1^ hr^-1^) and the number of reactions carrying flux in evolved *i*RH830 (blue triangle) and evolved *i*RH830 (yellow circle) models. The evolved solutions produce comparable biomass outputs. Ten evolved solutions are given for each, some duplicates are present.

Full evolved models with fluxes can be found in Supplementary File 3. Of the 383 unique reactions that carry flux in the evolved *i*RH830 models, 289 (75.5%) are found in all ten. For the *i*LF517 solutions, 301 of the 316 (95.3%) unique reactions that carry flux are found in all ten. This suggests that the smaller *S. glossinidius* model has fewer viable trajectories compared to the larger *S. praecaptivus* model. Of the 441 unique reactions across the 20 evolved models, 225 (51%) were found in all of the *i*RH830 and *i*LF517 evolved solutions. The biomass outputs for the evolved *i*RH830 and *i*LF517 solutions range from 0.064 to 0.281 (gr DW (mmol glucose)^-1^ hr^-1^), and 0.075 to 0.331 (gr DW (mmol glucose)^-1^ hr^-1^), respectively (Figure 7B).

## 4 Discussion

Classical studies of microbial evolution, whilst useful, are ultimately limited by their inherent inability to replicate adaptations over large evolutionary time scales. Here, we present a computational approach by combining a MOEA with FBA. We present the *Sodalis* system as a model for this. The *Sodalis* genus is ideal for the study of the evolution of symbiosis in this way, as within the genus are a free-living and a host restricted species, both well defined with complete genome sequences and existing protocols for culture.

This prototrophic model, *i*RH830, can produce a positive biomass output in both nutrient limited and nutrient rich conditions. As expected of a free-living organism, *i*RH830 exhibits a large amount of metabolic redundancy. We demonstrate the importance of thorough testing of FBA models, as described previously (Hall et al., 2019). Only through a large-scale phenotypic screen of carbon usage was the ability of *S. praecaptivus* to grow on xylitol alone identified. This had not been included in early iterations of *i*RH830 as xylitol is a carbon source rarely present in FBA models of Gram negative bacteria. It is possible that the unusual ability of *S. praecaptivus* to metabolise this important plant-derived sugar is related to the frequent presence of the *Sodalis* genus as a symbiont amongst insects.

The work here demonstrates the power of evolutionary algorithms in the study of symbiont evolution. A strength of this system is that removal of a reaction from the model is not irreversible; it is possible for a reaction to be added back into an individual at any point. Whilst there is no evidence currently for horizontal gene transfer within *S. praecaptivus*, the NSGA-II algorithm is only intended to be used as a tool to explore the possible evolutionary space rather than as a biologically accurate model of genome reduction. Previous examples of evolving minimal metabolic networks do not allow for full exploration of the possible evolutionary space (Pál et al., 2006). Decisions that are made at the start of process persist which, whilst biologically relevant, do not allow the full complement of evolutionary routes to be examined. It is not possible to know at what point during the transition to symbiosis the genes in *S. glossinidius* became pseudogenised. It is therefore valid, using the technique described here, to not attach significance to the temporal sequence in which the mutations occur. Allowing reactions to be added back into the model may also enable horizontal gene transfer to be incorporated, although whether this is relevant for the *Sodalis* genus remains to be established.

Supplying the ancestral *i*RH830 with contrasting growth media demonstrates the effect that nutrient provisioning may have on evolutionary trajectories of symbiotic bacteria. Exposure to famine reflects what might be expected in a nutrient-limited environment *in vivo*, in which evolutionary pressures result in the retention of pathways to synthesise key, essential metabolites. Here, this has shown to be particularly evident in the pathways retained for glycolysis/gluconeogenesis, the pentose phosphate pathway, and amino acid metabolism. This indicates that key pathways in central metabolism and for the synthesis of biomass components are being retained when the external environment is nutrient limited. The evolved solutions therefore reflect what is observed in symbiotic bacteria; the retention or loss of pathways can be used to inform about the microenvironment it resides within.

It has been shown that the evolved famine solutions contain a much greater number of core non-essential reactions that are present in a small percentage of the solutions. This suggests a lack of flexibility in the evolved network; either the reaction is found repeatedly, or not at all. This is not observed in the solutions provided with rich media (blood or sap), where a greater degree of flexibility is demonstrated by more reactions being included repeatedly across the evolutionary space. This implies that, *in vivo*, there are many possible trajectories for an early symbiont if there are sufficient nutrients in the microenvironment.

This tool can produce biologically relevant simulations that accurately reflect the metabolic pressures that symbionts are exposed to. An example of this was demonstrated by the investigation of key knockouts in *S. glossinidius*. The symbiont has a pseudogenisation in *ppc*, a key gene in central metabolism (Hall et al., 2019). It is not possible to deduce when this loss occurred from the genome annotation alone. By removing the PPC reaction from *S. praecaptivus* at the start of the *in silico* evolution, the resulting trajectories can be analysed and compared to WT. The loss of PPC appeared to have minimal effect on the resulting evolved populations compared to WT, in contrast to what was observed when the ASPTA reaction was removed at the start (Figure 4). This would indicate that, *in vivo*, the loss of the gene encoding the ASPTA reaction would have a greater impact on a bacterial symbiont if it was lost early in the relationship in comparison to the lower burden that the loss of the genes encoding PDH or PPC would have. This is interesting when considered in the context of the tsetse-*S. glossinidius* symbiosis. *S. glossinidius* has lost the *ppc* gene, whereas it has retained the genes encoding the PDH (*SG0467-9*) and ASPTA (*SG1006*) reactions (Dale and Maudlin, 1999; Hall et al., 2019). As the profile of ΔPPC evolution is similar to that of WT, it could be suggested that the *ppc* gene could have been lost early in evolutionary time without heavily bottlenecking *S. glossinidius* evolution subsequently. The gene encoding the ASPTA reaction may have been retained by *S. glossinidius* because of the detrimental impact that its loss may have caused. This is therefore a useful tool for making general predictions about when key pseudogenisations in insect-bacterial symbioses occurred.

It has been shown here that is it possible for *S. glossinidius* to reduce its metabolic network to approximately half of the size that it is currently. This provides support for the published hypothesis that *S. glossinidius* is a recent acquisition by the tsetse (Dale and Maudlin, 1999). The number of reactions remains slightly higher in evolved *i*RH830 models compared to evolved *i*LF517 solutions, possibly due to difficulties in finding the minima from a larger starting point. *i*RH830 can however be reduced down to look phenotypically similar to *i*LF517 at the level of biomass output, but with differences at the individual reaction level. These results suggest therefore that the route that *S. glossinidius* has taken within the tsetse is perhaps just one of several possible routes. The differences also indicate that *S. praecaptivus* may not be the ancestor that initiated the tsetse-*S. glossinidius* symbiosis.

The *Sodalis* genus, with a well-characterised free-living organism and symbiont relative, is a useful system to investigate the evolution of symbiosis. Previous uses of metabolic models to simulate evolution have focused on *E. coli* as a proof of concept. The tool described here has augmented knowledge about the temporal loss of key genes in *S. glossinidius* central metabolism. It has also investigated how *S. glossinidius* may have adapted to fluctuations in nutrient availability over time. It is not limited to this system, however. Combining FBA with a MOEA could be used for any organism for which a well-annotated genome is available. It could be applied not only to the evolution of symbiosis but to the directed evolution of, for example, industrially relevant microorganisms or to the study of rapid genome evolution in pathogenic bacteria.

## 5 Materials and Methods

### 5.1 Bacterial strains, growth conditions and reagents

*S. praecaptivus* was obtained from DMSZ (Brunswick, Germany). Working stocks were established by incubating starter cultures on LB (Merck, Darmstadt, Germany) agar plates overnight at 37°C. A single colony was then sub-cultured on to a fresh LB plate and incubated overnight at 37°C. A single colony was selected with a sterile pipette tip and used for downstream experimentation as per Biolog, Inc (Hayward, CA, USA) manufacturer protocol. Briefly, the colony was vortexed in IF-0 media before a redox dye was added (Biolog). Phenotypic microplates were used to screen for the ability of *S. praecaptivus* to grow on a range of carbon sources, using PM1 and PM2A microplates (Biolog). A 100 *µ*L bacterial suspension in the relevant media was added per well. Optical density was measured at 590 nm and 730 nm in a microplate reader (Epoch, BioTek, Winooski, VT, USA), and incubated with double orbital shaking at 37°C for 24 hours.

Discrepancies between *in silico* and *in vitro* Biolog results were reexamined by establishing individual cultures of *S. praecaptivus* in M9 salts in 96-well microplates, and supplemented with the metabolite of interest at a range of concentrations from 25 mM to 50 *µ*M. Cultures were incubated in a microplate reader with double orbital shaking at 37°C for 36 hours.

### 5.2 Construction of the *S. praecaptivus* metabolic network

The annotated *S. praecaptivus* genome sequence, CP006569.1, was downloaded from NCBI in GenBank format. Genes in *S. praecaptivus* with the same annotation as genes in the *E. coli* str. K-12 substr. MG1655 genome (ASM584v2) were highlighted, and the reactions encoded by these genes extracted from the BiGG Models database (King et al., 2016). These processes were automated using custom scripts written in Python.

FBA models of *S. glossinidius* (*i*LF517 (Hall et al., 2019)) and *E. coli* (*i*JO1366 (Orth et al., 2011; Orth and Palsson, 2012), *i*JR904 (Reed et al., 2003), *i*AF1260 (Feist et al., 2007)) were then used to aid the identification of missing reactions. The reactions and corresponding gene assignments in these published models were compared to the draft *S. praecaptivus* model. These gene assignments were then used to guide translated nucleotide and protein BLAST searches of the *S. praecaptivus* genome. KEGG (Kanehisa and Goto, 2000; Kanehisa et al., 2019) and EcoCyc (Keseler et al., 2017) databases were used to confirm the identity of the *E. coli* genes encoding each reaction. *S. glossinidius* gene assignments were taken from *i*LF517 (Hall et al., 2019). These orthologues in *E. coli* and *S. glossinidius*, with sequences taken from UniProt (Bateman et al., 2019), were used as BLAST search queries.

KEGG, BiGG Models, and MetaCyc (Caspi et al., 2007) were used to assign reaction stoichiometry. Candidate pseudogenes were aligned with known functional orthologues using ClustalX 2.1 (Larkin et al., 2007). Those with sequences missing or mutations in key residues were not included in the model. FBA and literature searches were used to identify and fill gaps in metabolic pathways appropriately (Thiele and Palsson, 2010).

### 5.3 Flux balance analysis

FBA solutions were generated using the GNA linear programming kit (GLPK) integrated with custom software in Java. Oxygen uptake was constrained to 20 mmol gr DW^-1^ hr^-1^, comparable to other models of free-living Gram negative bacteria. The uptake of ammonia, water, phosphate, sulphate, potassium, sodium, calcium, carbon dioxide, protons and essential transition metals was unconstrained for all media conditions. Cofactor constraints were implemented by introducing these metabolites to the biomass functions at small fluxes (1 × 10^−5^ mmol gr DW^-1^ hr^-1^) (Thomas et al., 2009). *i*RH830 was supplied with either 6 mmol gr DW^-1^ hr^-1^ GlcNAc and 1 mmol gr DW^-1^ hr^-1^ thiamine (henceforth “famine”), a tsetse-specific media (henceforth “blood”, Table S1), or a sap-inspired media (from Krishnan et al. (2011), henceforth “sap”). Full recipes are provided in Supplementary File 1. The phenotype was considered viable if the biomass production rate was greater than 1 x 10^−4^ gr DW (mmol glucose)^-1^ hr^-1^. Futile cycles, closed loops of a number of reactions, were detected by the presence of unsustainably large fluxes. Futile cycles often occur when several reversible reactions are present in which the product of one becomes the substrate of another. These reactions were examined individually, and solved by adjusting the reversability with guidance from EcoCyc and BiGG Models.

To investigate the concordance between the *in vitro* screen and the *in silico* outputs, *i*RH830 was, where possible, supplemented with the carbon sources analysed at an exogenous concentration of 6 mmol gr DW^-1^ hr^-1^. A qualitative presence/absence of a positive biomass output was noted. Full description of the model is provided in Supplementary File 1.

### 5.4 Robustness analysis

Robustness analysis of the *i*RH830 network was executed using COBRApy (Ebrahim et al., 2013) to conduct single reaction deletions. *i*RH830 was supplied with either famine or blood media under aerated conditions. The flux through reactions was set to zero individually and the resulting effect on biomass output measured. Reactions were categorised as essential if the resulting biomass output was less than 1 x 10^−3^ gr DW (mmol glucose)^-1^ hr^-1^.

### 5.5 Implementation of multi-objective evolutionary algorithm

A MOEA was used to explore possible evolutionary trajectories in the *Sodalis* genus. An overview of the process is provided in Figure 8. The non-dominated sorted genetic algorithm (NSGA-II) (Deb et al., 2002) from the Distributed Evolutionary Algorithms in Python (DEAP) (De Rainville et al., 2012) package was used in combination with the COBRApy package (Ebrahim et al., 2013) for FBA evaluation. Equal weight was placed on reducing the number of reactions used in the model whilst maximising the biomass output.

**Figure 8:**
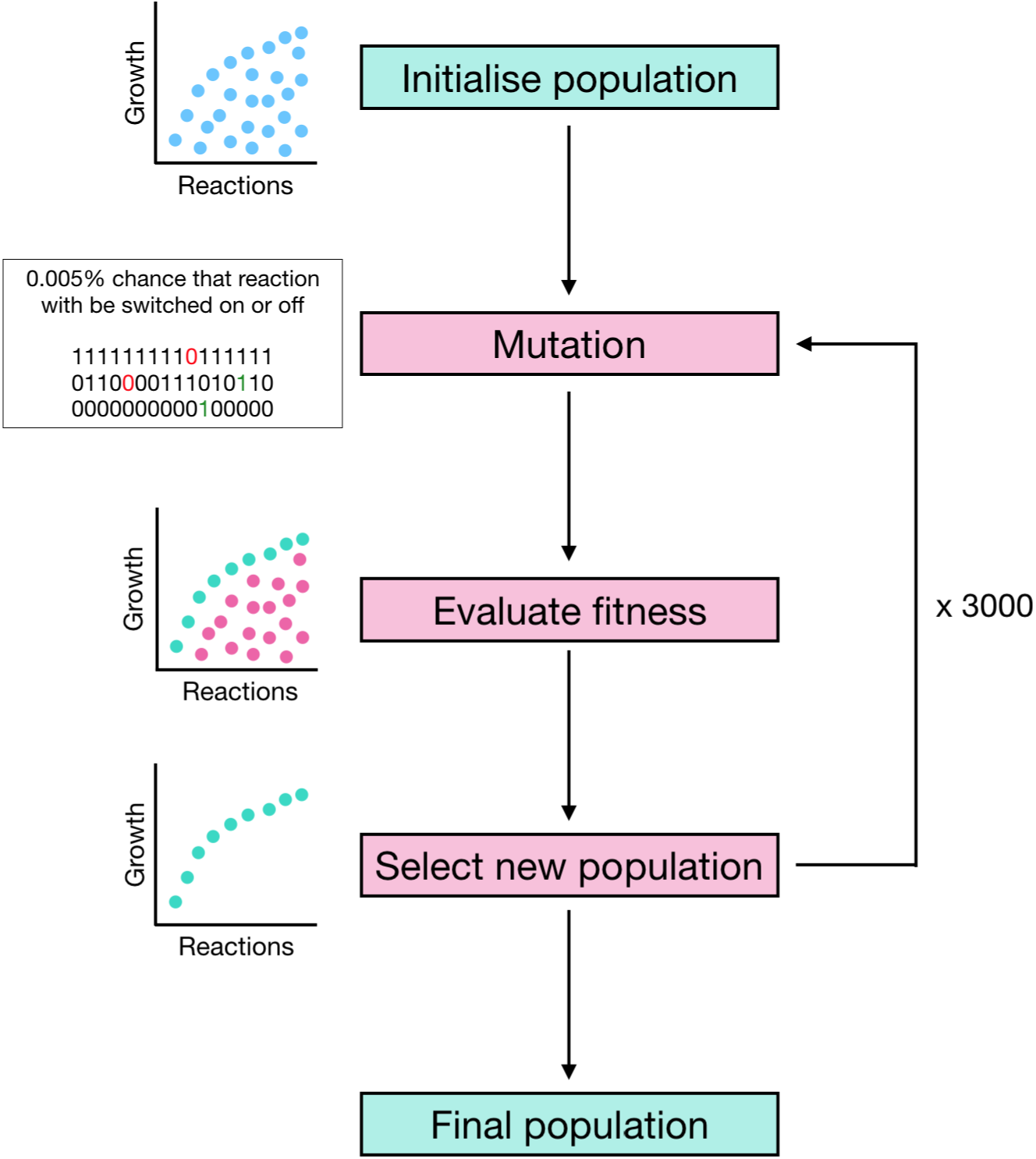
Process of the MOEA. A starting population of individuals is initialised, and the fitness calculated by solving the FBA model to calculate biomass output and the number of active reactions. The population is then copied, allowed to mutate and the fitness evaluated again. A new population is selected from the original and copy populations. Green boxes represent the start and final populations, pink boxes represent the iterative process of mutation and selection.

#### 5.5.1 Population initiation

Prior to starting an evolutionary run, reactions essential to growth were identified using a single reaction knockout. Essential reactions were defined as those producing a biomass output of less than 1 × 10^−3^ gr DW (mmol glucose)^-1^ hr^-1^. Reactions that were identified as essential were not included in the subsequent mutation strategy, therefore reducing the solution space and computational time taken to run the MOEA. The essential reactions were added back to the evolved populations for downstream analysis.

A population of 100 genotype clones, where all non-essential reactions are active, was created (Figure 8). Each genotype consisted of a binary number, where a 1 or 0 corresponded to the reaction being active or inactive, respectively. This is a proxy for gene loss, where a one-to-one gene-protein-reaction mapping is assumed. All post-evolution analysis focused on the reactions lost or retained.

#### 5.5.2 Mutation

Mutation was performed on each genotype by flipping the value of each reaction with a probability of 0.005 (Figure 8). The fitness of each individual is evaluated by solving the FBA model to calculate both its biomass output and the sum of number of active reactions.

#### 5.5.3 Fitness evaluation and selection

The population was first evaluated for non-dominated individuals. This gave a population of individuals that has the highest biomass output for their current number of active reactions (Figure 8). From the non-dominated population, the Euclidean distance between each individual was calculated. A greater priority was given to selecting individuals with a larger Euclidean distance. This prevented the clustering of similar potential solutions, thereby reducing the likelihood of becoming trapped in sub-optimal local minima within the search space. The resulting population maximised the convergence on the highest biomass output, lowest number of reactions, and the distribution of those solutions. There will be a set of solutions whereby the number of reactions cannot be minimised further without also reducing the corresponding biomass output. This set of solutions is known as a Pareto front. The algorithm was repeated for 3000 generations to produce genotypes that converged. This indicated that minimal new solutions were being found. The biomass output from the slim optimisation COBRApy function and the summation of the number of active reactions was used to evaluate the fitness.

#### 5.5.4 MOEA variations

The MOEA was run under several conditions in order to investigate aspects of symbiont evolution. Full details are provided in Table 3. Scenario iinvestigated the trajectories taken when the *S. praecaptivus* model was provided with blood, sap, and famine growth media. In Scenario ii, gene knockouts were simulated by removing individual reactions from the *S. praecaptivus* model prior to commencing the evolution. The reactions chosen were ASPTA, PDH, and PPC. In Scenario iii, the MOEA was applied to a model of *S. glossinidius* metabolism, *i*LF517 (Hall et al., 2019). Here, *i*LF517 was supplied with the blood medium for 3000 generations.

#### 5.5.5 Analysis of evolved populations

To identify key reactions in the evolved populations, individuals were selected from each condition and the remaining non-essential reactions extracted. The subset of reactions that were present in every individual selected were designated as “core non-essentials”, and will be referred to hereafter as such. When examining the similarity between evolved models, exchange reactions and reactions carrying zero flux were discounted.

## 6 Acknowledgements

RJH was funded by the BBSRC White Rose DTP (BB/M011151/1) and ST by the Wellcome Trust CIDCATS programme (WT095024MA).

## 8 Supplementary

**Figure S1:**
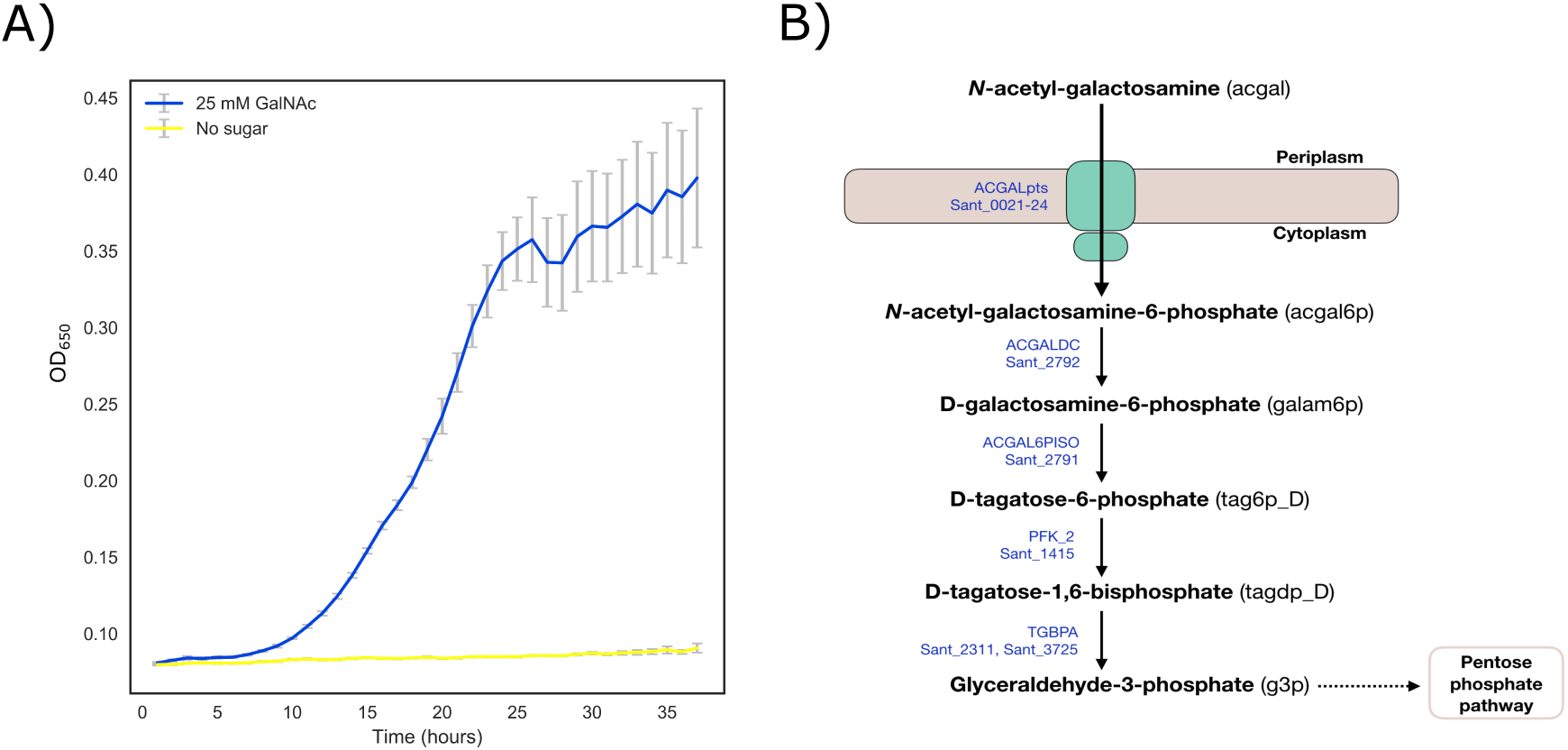
GalNAc use by *S. praecaptivus*. (**A)** Growth of *S. praecaptivus* on 25 mM GalNAc (blue) and in M9 salts with no added sugar (yellow) over 36 hours. Measurements in sextuplicate, error bars SEM. (**B**) The putative GalNAc degradation pathway in *S. praecaptivus*. Reaction names and *S. praecaptivus* gene assignments are given in blue.

**Table S1:** *In silico* blood medium for *i*RH830. The maximum permitted and actual flux values for each metabolite (mmol gr DW^-1^ hr^-1^) is presented.

| Metabolite | Name | Flux (permitted) | Flux (actual) |
| --- | --- | --- | --- |
| acgam | GlcNAc | 6 | 6 |
| ala_L | L-alanine | 3.41 | 3.41 |
| arg_L | L-arginine | 1.54 | 1.14 |
| asn_L | L-asparagine | 0.58 | 0.58 |
| asp_L | L-aspartate | 0.03 | 0.03 |
| cys_L | L-cysteine | 1.18 | 0 |
| gln_L | L-glutamine | 8.3 | 0 |
| glu_L | L-glutamate | 0.7 | 0.7 |
| gly | Glycine | 1.54 | 1.54 |
| his_L | L-histidine | 1.15 | 0.11 |
| ile_L | L-isoleucine | 0.89 | 0.33 |
| leu_L | L-leucine | 1.69 | 0.52 |
| lys_L | L-lysine | 2.72 | 0.39 |
| met_L | L-methionine | 0.38 | 0.18 |
| orn | Ornithine | 0.72 | 0 |
| phe_L | L-phenylalanine | 0.84 | 0.21 |
| pro_L | L-proline | 2.36 | 2.36 |
| ser_L | L-serine | 1.12 | 1.12 |
| taur | Taurine | 0.55 | 0 |
| thm | Thiamine | 1 | 0.00001 |
| thr_L | L-threonine | 1.39 | 0 |
| trp_L | L-tryptophan | 1.11 | 0.07 |
| tyr_L | L-tyrosine | 1.03 | 0.16 |
| val_L | L-valine | 2.88 | 0.49 |

**Table S2:**
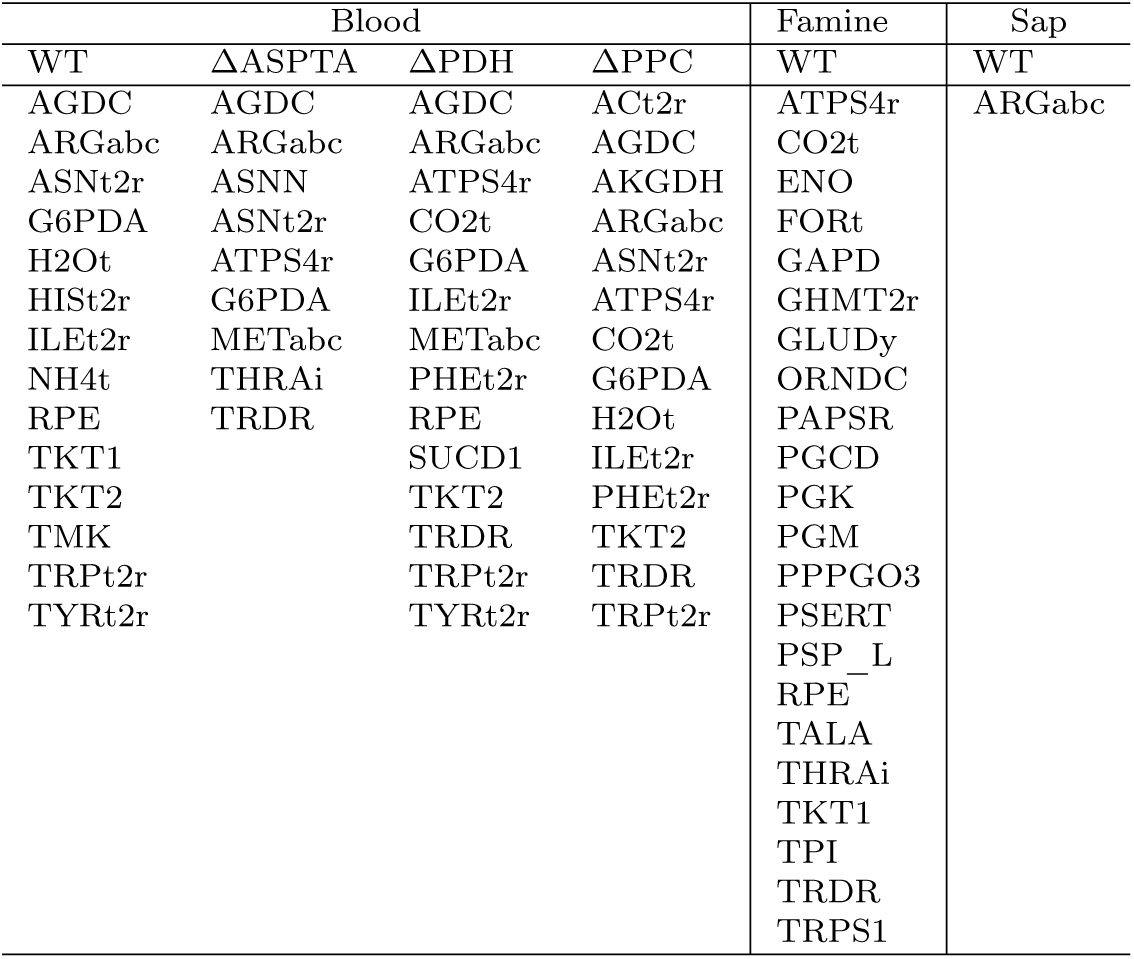
Core non-essential reactions. Reactions conserved between the samples of evolved *i*RH830 models under each condition.

